# Genetic Mechanisms underlying Preference-Performance Mismatches: Insights from a Specialized Native Herbivore on an Invasive Toxic Plant

**DOI:** 10.1101/2024.02.14.580222

**Authors:** Nitin Ravikanthachari, Carol L Boggs

**Author notes:** Division of Biological Sciences, University of Montana, Missoula, Montana (Nitin Ravikanthachari).

## Abstract

Specialist phytophagous insects have a narrow hostplant range for optimal development and survival. Mismatches between female oviposition preference and larval performance can lead to high fitness costs. Understanding the mechanistic basis of this decoupling can help us understand evolutionary constraints and aid in predicting outcomes of error-prone oviposition. We investigated the causes for preference-performance mismatches in a specialist native herbivore laying eggs on an invasive toxic plant. Transcriptomic analyses revealed no differential gene expression in gustatory/olfactory organs of adult females with different oviposition preference, but larvae exhibit host-plant-specific gene expression signatures. The larvae feeding on toxic plants showed lower expression of specialized detoxification enzymes and higher expression of general digestive enzymes, indicating the inability of larvae to detoxify toxic compounds present in the toxic plants. We additionally found that genes related to successful detoxification and adaptive feeding were enriched in larvae feeding on native plants, while genes related to toxic responses, apoptosis and accelerated development were enriched in larvae feeding on toxic plants. Our findings dissect genetic mechanisms behind preference-performance mismatches, quantifying the impact of error-prone oviposition on larval performance in specialized species interaction.

## Introduction

Herbivorous insects and the plants they feed on are a classic example of co-evolution [1–3]. Herbivorous insects have evolved to feed on a narrow range of hostplants by adapting to circumvent specialized plant defenses [4–6]. In holometabolous insects such as Lepidoptera, where the larval stages are relatively immobile, the success of the herbivore depends on the female choosing a suitable host that maximizes the survival of the resulting offspring [7–9]. This results in strong selection for female oviposition preference. This idea, known as preference-performance hypothesis, postulates that the preference of a hostplant by the female is positively correlated with performance of the larvae on that hostplant [10,11]. Female host choice is dependent on a multitude of factors including availability of suitable hosts, hostplant specialization, hostplant density and presence of natural enemies [12–14]. Support for the preference-performance hypothesis ranges from none to strong correlation in various insect groups [11,15,16]. A meta-analysis showed that oligophagous insects (insects feeding on several plant genera within a plant family) had a stronger correlation compared to polyphagous insects (insects feeding on several plant families) [17]. However, for specialized insects, the evidence is mixed. One underlying hypothesis for this mixed evidence is that cue similarity between hosts and non-hosts often leads to error prone oviposition in females [18,19].

Error prone oviposition can result in the following scenarios [20]. It can be adaptive if the larvae can survive on the novel hostplant, thus leading to an increase in the diet breadth of the herbivore and adaptation to new hostplants [21,22]. In fact, error prone oviposition has often been cited as the fuel driving hostplant range in Lepidoptera and has been documented in many species [21,23–25]. The success of the larvae on the novel host depends on underlying variation in detoxification enzymes, the degree of chemical similarity of the novel host to the hosts that the larvae have co-evolved with and the evolutionary history of host use in the insect species [26–29].

Alternatively, error prone oviposition can lead to high fitness costs if the larvae do not possess behavioral or physiological mechanisms to utilize the novel host plant, thus leading to higher mortality and maladaptation [19,30–33]. Repeated error prone oviposition can potentially lead to the larvae evolving the ability to feed on the plant provided that new mutations or underlying standing genetic variation leads to evolution of key innovations in detoxification machinery [23,34].

In certain scenarios, known as evolutionary traps, abrupt or rapid environmental change can lead to instances where evolved reliable cues fail, resulting in repeated oviposition error [19,30,31,35,36]. This could occur either due to female’s inability to distinguish between hosts and non-hosts to avoid costly mistakes or the failure of larvae to incorporate the novel host due to behavioral or physiological constraints [31–33,36–38]. Identifying the underlying mechanisms that lead to decoupling cue-response systems as well as preference-performance can shed light on the constraints of evolution in species interactions.

Although evolutionary traps have been documented in many systems, mechanistic processes underlying persistent evolutionary traps are not well understood [39–42]. Here we use a system of a specialized native herbivore that oviposits on a lethal, invasive plant that causes high mortality in the larvae to dissect the underlying genetic mechanisms affecting adult female oviposition, and larval feeding on the plant. We employ a transcriptomic approach to quantify differential gene expression to address the hypotheses that a) females preferring the toxic lethal plant differ in their expression repertoire of sensory/gustatory genes compared to those preferring the native plant and b) larvae that feed on the toxic plant exhibit transcriptomic signatures of impaired feeding and toxicity (upregulation of stress responses) compared to those feeding on the native host plant. Our study adds to the extensive literature on preference-performance studies by elucidating the underlying genetic mechanisms that decouple female preference and larval performance.

## Methods

### Study organisms

*Pieris macdunnoughii* Remington, 1954 (Pieridae; formerly *P. napi macdunnoughii*) [43] is a montane butterfly distributed in the Southern Rocky Mountains in North America and is a specialist herbivore: the females lay eggs on and the larvae feed on native Brassicaceae [37,44–46]. *Pieris macdunnoughii*, like other species in the *Pieris* species complex, has evolved detoxification enzymes to overcome the toxicity of glucosinolates (secondary metabolites) in Brassicaceae [34,47]. However, larvae experience high mortality when they encounter novel mustards whose glucosinolates/secondary metabolites differ from those with which they have locally co-evolved [31,33,44,48,49].

*Thlaspi arvense* (L.) (Brassicaceae) is a plant native to temperate Eurasia that was introduced to North America in the 1800s [37]. It was introduced to Gunnison County between 1850s to 1870s with the earliest herbarium record dating back to 1929 when the Rocky Mountain Laboratory’s herbarium was established. Thus, *T. arvense* has been present in the habitat for at least 94 years [37]. *Thlaspi arvense* is an early successional plant which colonizes disturbed soil and is known to occur up to 2900m [50,51].

The glucosinolate profile of *T*. *arvense* is comprised mainly of the aliphatic glucosinolate sinigrin, in contrast to native mustards that contain both aliphatic and aromatic glucosinolates [52]. *Pieris macdunnoughii* females recognize *T. arvense* as a potential host plant in areas where both co-occur due to cue similarity with the native plants [37,46]. Co-occurrence of *T. arvense* with other native mustards has been estimated to cause a fitness loss to females of 3% due to larval survival [46,53]. Therefore, there is strong selection on females of *P. macdunnoughii* to avoid laying eggs on the plant and for escape from the evolutionary trap.

### Oviposition choice experiment

Gravid females were collected in the East River Valley, Gunnison County, Colorado (38.9664° N, 106.9896° W, 2,896 m a.s.l) in 2019 using an aerial net. In the lab, the females were fed twice a day with 25% honey-water solution. Females were housed in plastic cages in an environmental chamber at 27°C during the day and at 18°C at night on an 18:6 L:D cycle. The females were provided with one rooted whole plant each of *T. arvense* and *Cardamine cordifolia* A. Gray (Brassicaceae), a primary native hostplant. The larval host plants were visually matched by approximate leaf size and by plant phenology (pre-flowering stage). The females were allowed to lay eggs on the host plants and the eggs from each plant were counted every morning. Preference for a hostplant was quantified as follows: preference for *T. arvense* if >80% of eggs laid on *T. arvense*; preference for *C. cordifolia* if >80% of eggs laid on *C. cordifolia* and equal preference if < 80% eggs laid on either plant. The females were removed while actively ovipositing once they had laid a total of fifty eggs, and their forelegs, antennae and head were stored in RNAlater at -20°C.

### Larval feeding assays

The larvae from eggs laid by females in the oviposition choice trails were reared in an environmental chamber under the same conditions used during female oviposition experiment. The larvae were reared on young leaves of *Raphanus sativus* L. (Brassicaceae) until they reached third instar. Five third instar larvae from each female were provided either with a potted whole plant of *C. cordifolia* or *T. arvense* after 12 hours of starvation. The larvae were allowed to feed for 24 hours. After 24 hours, the larvae were visually examined to confirm that their midguts were at least partially filled with plant material. The larvae were then dissected in PBS solution, degutted, plant material removed, and their mouth parts and guts were stored in RNAlater.

### RNA extraction and sequencing

Total RNA was extracted from 10 females (head, antennae, and forelegs) each preferring *T. arvense* and *C. cordifolia* and from 10 third instar larvae (gut and mouthparts) each fed on *C. cordifolia* and *T. arvense* using a QIAGEN RNeasy kit following manufacturer’s protocol. Total RNA from all samples were sent to MedGenome for cDNA library preparation and RNA sequencing. cDNA library preparation was carried out using Illumina TruSeq stranded mRNA kit followed by 20M paired reads (40M total) sequencing on a Novaseq S4 platform.

### Read mapping

Demultiplexed raw Illumina reads were checked for quality using FastQC [54]. Reads were trimmed using Trimmomatic v. 0.39 [55] with the following options: Sliding window: 4:20, minimum length: 25, and adapter sequences were cleaned using the option TruSeq3. Cleaned transcripts were mapped to the reference genome [56] using genome guided STAR assembly [57,58]. We first generated genome indices and the genome directory, followed by mapping reads to the genome with default options. 94% of the reads were correctly mapped to the genome. The resulting BAM files were sorted by genome coordinates using SAMtools [59] sort and indexed using SAMtools index. Mapped reads were then quantified using featureCounts (useMetaFeatures geneid) [60] and ran through Rsubread [61] in the R statistical environment v. 4.0 [62]. We ran featureCounts separately on adult female and larval mapped reads to obtain their respective expression profiles.

### Differential gene expression analysis

Differential gene expression (DGE) analyses for adult females and larvae were carried out separately in edgeR [63]. We used CPM-TMM log2 transformation for further filtering our dataset. The raw counts were filtered based on abundance and then normalized using count-per-million (CPM) to account for differences in library sizes among samples in edgeR. We additionally used trimmed mean of M values (TMM) for cross-sample normalization using the option “calcNormFactors” in edgeR. Genes that were lowly expressed were filtered using the HTSFilter followed by differential gene expression analysis using the Fisher’s exact test approach in edgeR. The differentially expressed genes were further filtered using a false discovery rate of 1% using the BH correction method. We set a cut-off of a minimal fold change (FC) of 1.5 between treatments and an FDR of p-value <1e-3 for assessing significant differential expression. The cut-off values for FC and FDR were set based on prior studies to increase the signal to noise ratio and to reduce non hostplant specific patterns [64,65]. We used the hclust function in R package stats for hierarchical clustering of significantly differentially expressed transcripts with a k value of 3 based on results from the sum of square means (SI fig 1). We additionally visualized differentially expressed genes using a heatmap using the pheatmap package in R [66].

**Figure 1:**
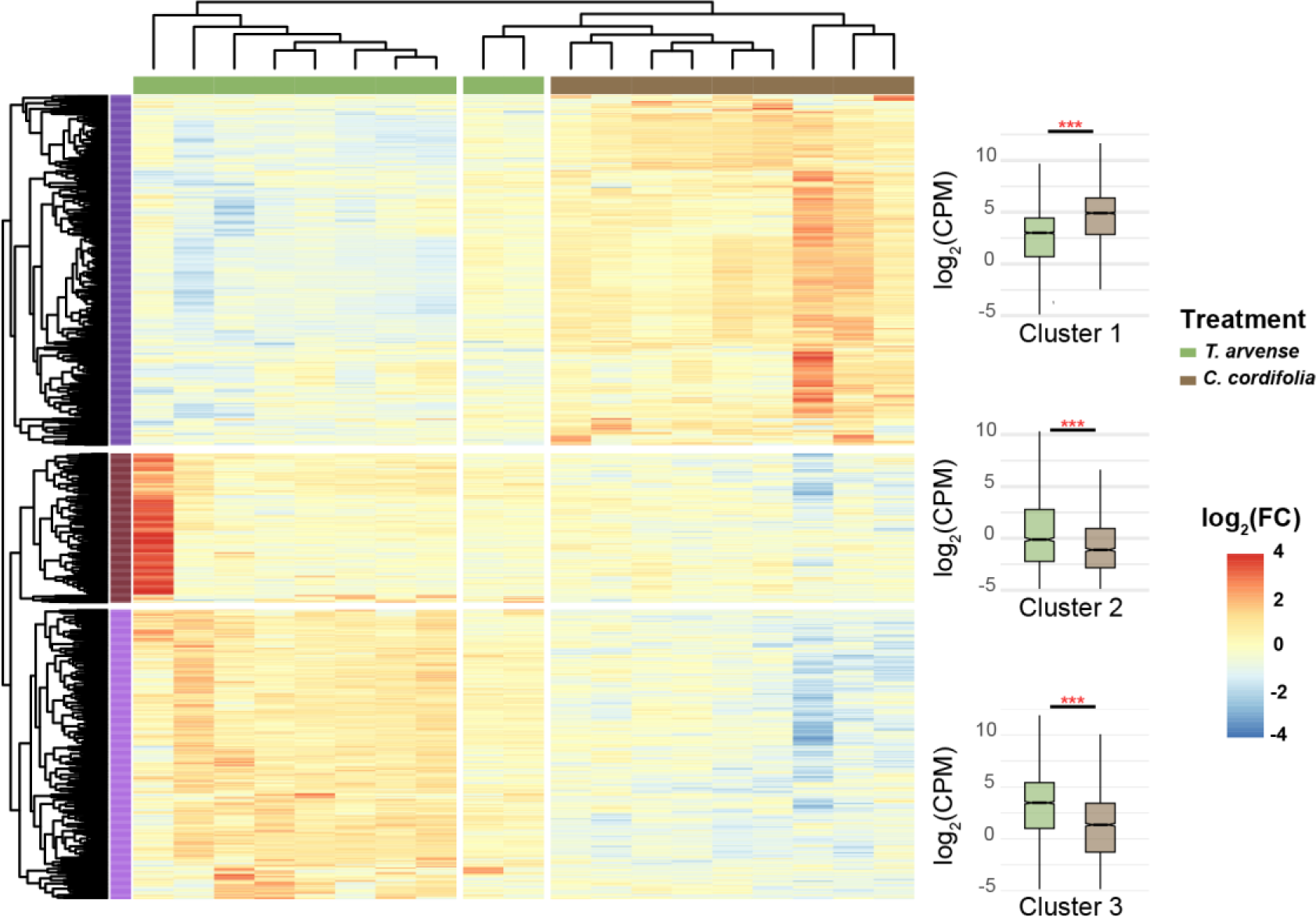
a) Heatmap showing log (FC) in larvae in response to feeding on *C. cordifolia* (brown) and *T. arvense* (green). Individuals (columns) and transcripts (rows) are arranged by hierarchical clustering of expression profiles. b) Individual gene expression profile corresponding to each cluster of gene expression represented in panel a.

### Gene expression in plant compound detoxification genes

We obtained the identity of the genes that were differentially expressed by linking the gene names to their respective protein names in the protein family (Pfam) database [67]. We extracted genes that corresponded to the six gene families involved in detoxification of plant compounds (Cytochrome p450s, Carboxylesterase, Glutathione-S-Transferase, Glucuronosyltransferase, Trypsin, and insect cuticle protein) and grouped them to their respective clusters. We then ran a paired t-test to assess significance among the different genes involved in detoxification between the two treatments.

### Weighted gene co-expression analysis

We utilized the WGCNA package in R [68] to identify plant-specific co-expressed genes in the larval treatments using TMM normalized gene counts (see above for description of normalization). For network construction, we selected a soft clustering power of 6 based on a scale-free topology model fit (SI fig 2; r2>0.9). The network was built using signed networks with Pearson correlation, requiring a minimum of 40 genes per module with a deep split value of 3 and merge cut height of 0.3. Subsequently, we tested the network for module-trait correlations with plant treatment as a binary variable (*T. arvense* = 1, *C. cordifolia* = 0). We ran a linear fit model using the *lmfit* function of the limma package [69] to identify modules of interest (module-trait correlations that showed the highest difference between plant treatments).

**Figure 2:**
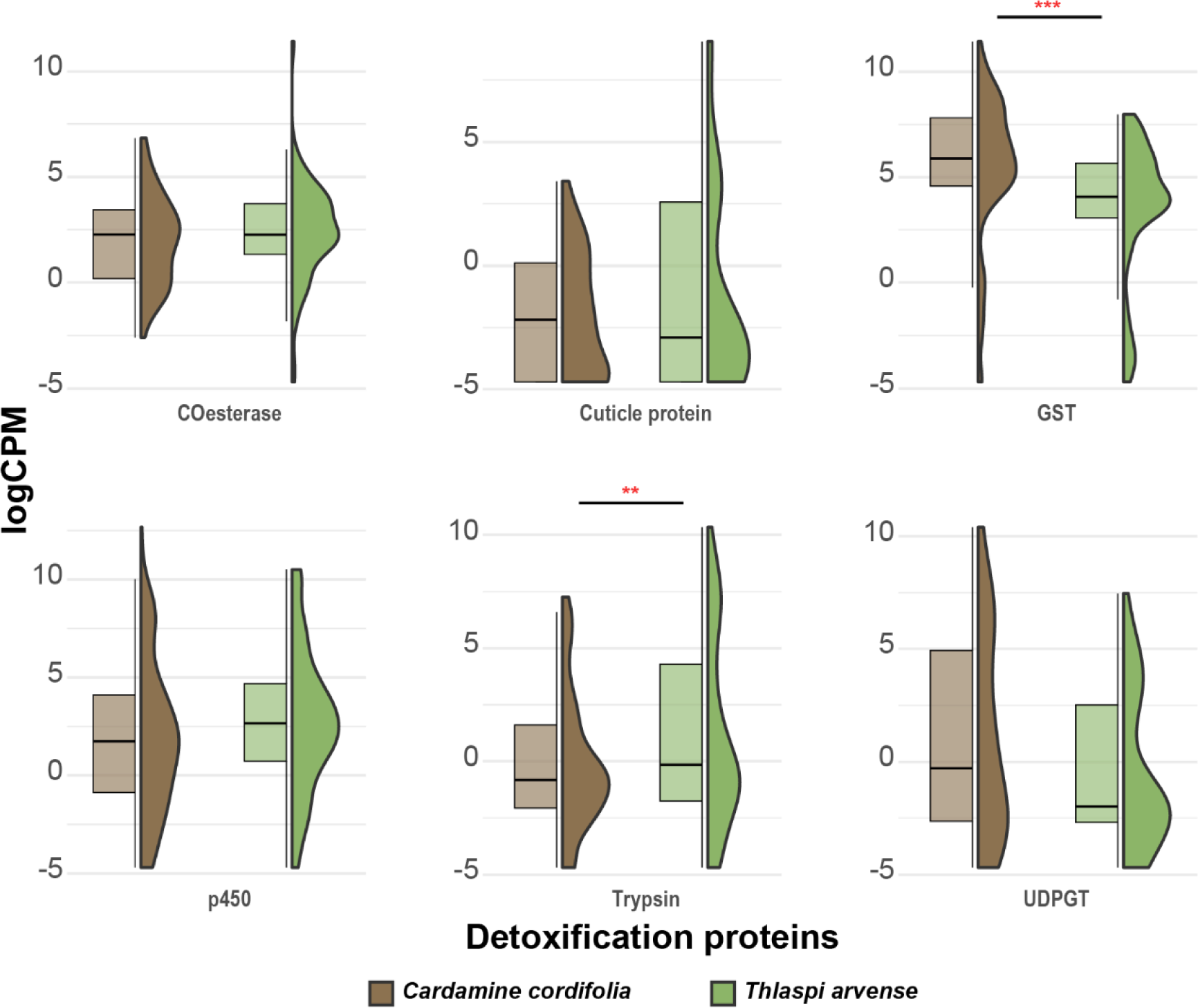
Gene expression of various plant compound detoxification genes grouped based on Pfam database annotation between larvae feeding on *T. arvense* and *C. cordifolia*.

We used two matrices to further identify specific genes associated with feeding on *T. arvense* from the modules of interest. First, we calculated gene significance and module membership for each gene in our dataset. We calculated gene significance for a gene as the correlation between the normalized gene expression of the gene and the treatment variable. We calculated module membership for each gene as the correlation between the normalized gene expression of the gene and the module eigengene values. Additionally, we calculated gene connectivity (a measure of node density for each gene) using the *signedKME* function in the WGCNA package. We considered genes as associated with *T. arvense* if they showed >0.6 correlation between gene significance vs module membership and scored >0.8 for gene connectivity based on author recommendations. Second, we used the *chooseTopHubsInEachModule* function in WGCNA to identify the top genes in the modules of interest. Hubs are central genes in each module that are highly connected.

### Gene set enrichment

We generated gene ontology (GO) annotation of the *P. macdunnoughii* genome using eggNOG-Mapper [70] by linking the annotated genes to their respective protein names using the protein family (Pfam) database to obtain GO terms. The GO annotation consisted of 15477 genes that were used as input for topGO [71] to quantify gene set enrichment in the differentially expressed and co-expressed genes. We used the parent-child algorithm with a two-tailed Fisher’s test to identify enriched GO terms based on biological processes (BP). We calculated gene set enrichment separately for a) each of the 3 distinct hierarchical gene clusters in the differentially expressed gene dataset and b) for filtered genes in each module of interest. We considered GO terms as significant for those that overlapped between the DGE dataset and WGCNA dataset.

The final list of enriched GO terms were then run through REVIGO [72] to cluster the terms and to reduce redundancy by identifying similarity between the GO terms.

## Results

### Variation in female oviposition preference

We tested 56 *Pieris macdunnoughii* females collected from the East River valley. 14 females preferred *Cardamine cordifolia*, and 10 females preferred *Thlaspi arvense*. Nine females had equal preference for both plants and 25 females did not meet the threshold of 50 eggs to quantify preference.

### No differential gene expression underlying female oviposition preference

Gene expression profiles of female *P. macdunnoughii* preferring *T. arvense* and *C. cordifolia* were similar. Our analysis consisting of females who laid 100% of their eggs on either of the plant (n=3/group, total 6 females) also did not identify any DE genes.

### Hostplant-specific changes in differential expression of genes underlying larval feeding

We found 1489 (FDR <0.01) overall differences in gene expression profiles between larvae that fed on *T. arvense* and *C. cordifolia*. Hierarchical clustering of individuals showed that replicates of larvae of each treatment were broadly similar to each other compared to those from the other treatments (fig. 1a), suggesting hostplant specific transcriptome patterns. However, two larvae that fed on *T. arvense* clustered with larval profiles from *C. cordifolia*. The hierarchical clustering of transcripts revealed 910 genes (log_2_FC >1.5 and FDR <0.01) that were differentially expressed between larvae that fed on *C. cordifolia* compared to *T. arvense*. These transcripts split into three distinct clusters (fig. 1a). Genes in clusters 1 (n=404 genes; SI table 1; paired t-test, p<0.001) were upregulated in larvae that fed on *C. cordifolia*. Genes in cluster 2 (n=172 genes; SI table 1; paired t-test, p<0.001) and 3 (n=334 genes; SI table 1; paired t-test, p<0.001) were upregulated in larvae that fed on *T. arvense*.

### Genes underlying detoxification of plant compounds differentially expressed between diet treatments

We found significantly higher expression of Glutathione-S-transferase (fig. 2; SI table 2; paired t-test, p<0.001) in larvae feeding on *C. cordifolia*. The larvae feeding on *T. arvense* had higher expression of trypsin (fig. 2; SI table 2; paired t-test, p=0.002). Glucuronosyltransferase (fig. 2; SI table 2; paired t-test, p=0.059) showed higher expression in larvae feeding on *C. cordifolia*, but the differences were marginally insignificant. Cytochrome p450s (fig. 2; SI table 2; paired t-test, p=0.09), Carboxylesterase (fig. 2; SI table 2; paired t-test, p=0.288) and cuticle proteins (fig. 2; SI table 2; paired t-test, p=0.288) showed no differences in their gene expression between the two treatments.

### Co-expression of genes indicate host plant specific functional gene modules

Our network analysis identified thirty-three modules (SI fig 3). Five of the thirty-three modules showed significant correlation with *T. arvense* (fig. 3). Three of those five modules (module blue, module black and module green) were positively correlated with *T. arvense* and two modules (module magenta and module turquoise) were negatively correlated with *T. arvense* (i.e. upregulated in *C. cordifolia*). We identified 2353 genes as significantly associated with *T. arvense* (gene significance vs module membership r^2^>0.6 and gene connectivity r^2^>0.8) across the five modules. 1795 genes showed a positive association with *T. arvense* (genes in modules blue, black, and green) and 1921 genes showed negative association with *T. arvense* (genes in modules magenta and turquoise).

**Figure 3:**
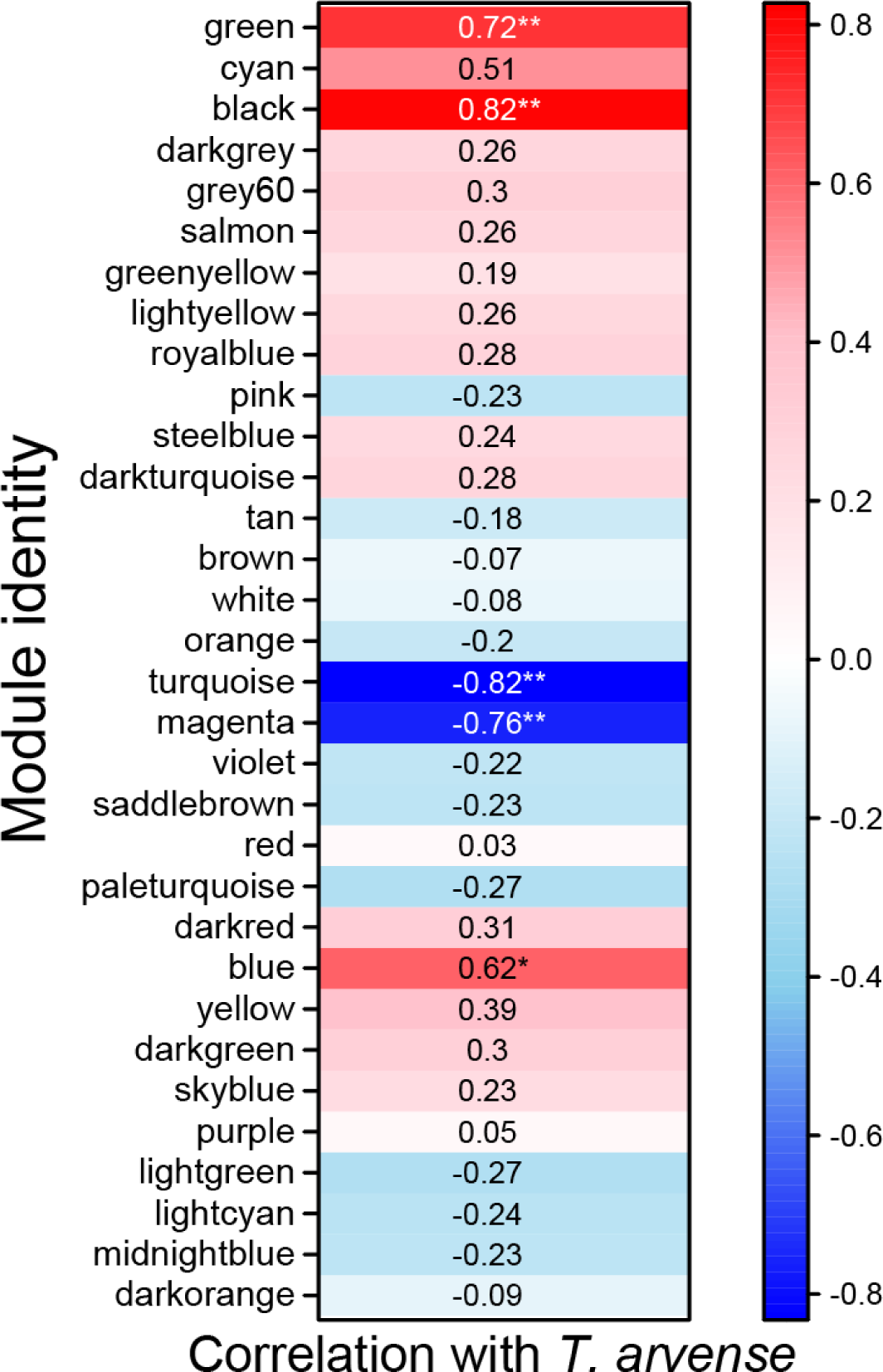
Module correlation with feeding on T. arvense. Modules represent clusters whose genes are co-expressed. Colors represent direction of correlation: red represents positive correlation of module with T. arvense and blue represents negative correlation with T. arvense (i.e. enriched in larvae feeding on C. cordifolia) Asterisk represents significant correlations.

**Figure 4:**
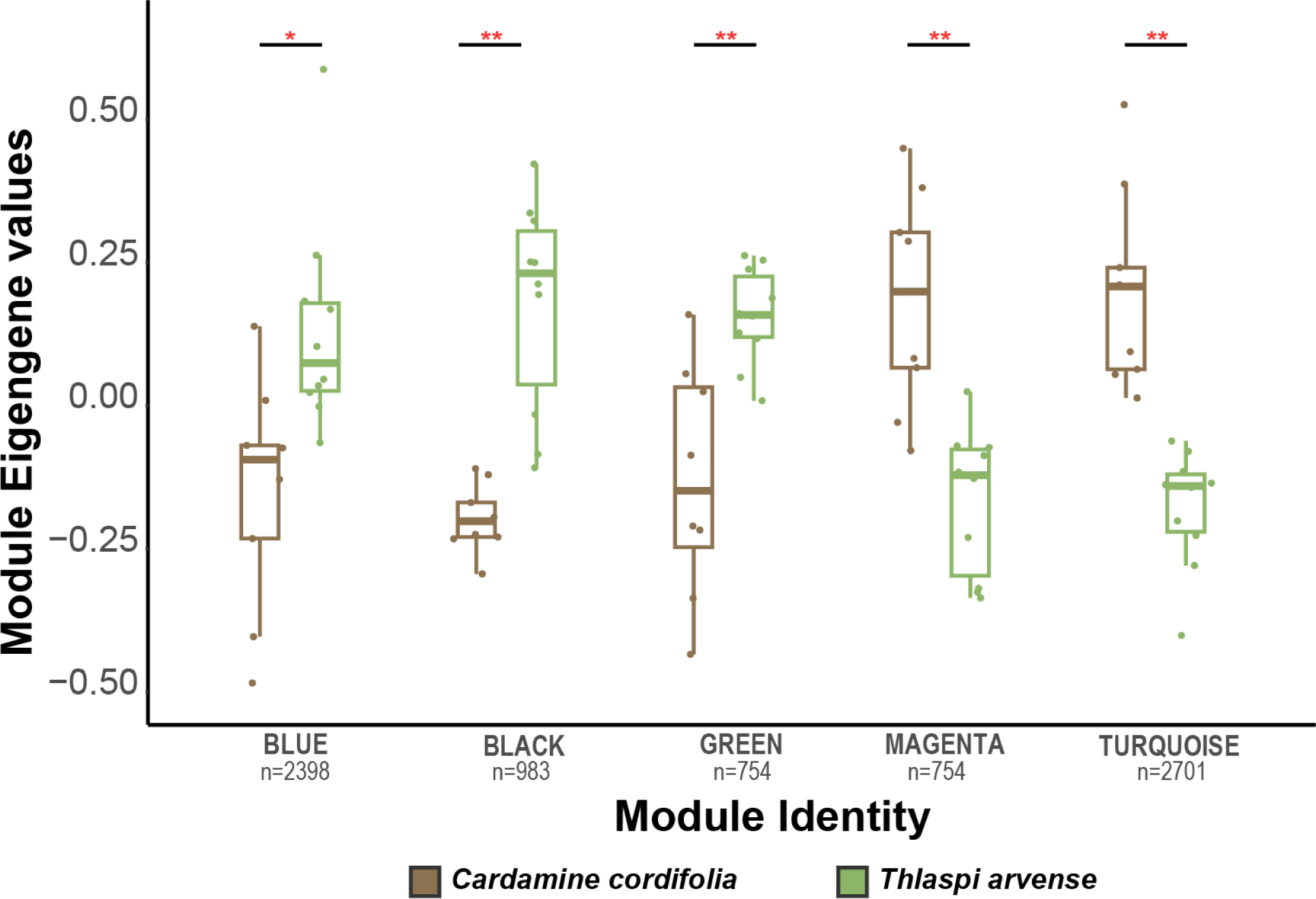
Differences in module eigengene values between treatments. Only modules that are show significant differences between treatments are shown.

Our analysis identified several hub genes of interest (implicated in metabolism, development, and/or immune process) in each treatment (SI table 3). Broadly, in modules that were positively correlated with feeding on *T. arvense*, genes related to lipid metabolism (Low density lipoprotein receptor-related protein 4 and C2 domain-containing protein), general metabolism (ABC transporter domain-containing protein, dipeptidase, ubiquitin hydrolase 1 and E3 ubiquitin-protein ligase), response to stress (Nicotinic acetylcholine receptor alpha 3 subunit, receptor protein serine/threonine kinase) and immune response (CUB domain-containing protein and EGF-like domain-containing protein) were identified as the top hub genes. In larvae feeding on *C. cordifolia*, genes related to DNA/RNA synthesis (WKF domain-containing protein, Exoribonuclease phosphorolytic domain-containing protein, Ribosomal protein L1, Menin and Tudor domain-containing protein) and carbohydrate metabolism (GH18 domain-containing protein) were the top hub genes.

### Gene set enrichment analysis reveals host-plant specific effects on critical larval traits

Overlap between co-expression modules and differential gene expression (clusters 2 and 3) in larvae feeding on *T. arvense* (SI table 4) showed that gene ontology terms involved in glucose metabolism (GO:0009052, GO:0019682, GO:0034641 and GO:0019682), lipid metabolism (GO:0051792), amino acid/nitrogen metabolism (GO:0000050, GO:0006546, GO:0034641, GO:1901564, GO:0043603, GO:0044271), protein metabolism (GO:0008213, GO:0009249, GO:0006457, GO:0006518, GO:0018195, GO:0070207, GO:0071826, GO:2000257, GO:0051205 and GO:0070585), nucleic acid metabolism (GO:0016072, GO:0046385 and GO:1901566), response to stress (GO:0009585 and GO:0006979), cell death and apoptosis (GO:0097190, GO:0034614, GO:0045476 and GO:0071478), immune response (GO:0001779, GO:0032695, GO:0030449, GO:0039531, GO:0043053, GO:0032814, GO:0070207 and GO:0002673) and development (GO:0071390, GO:0021697, GO:0061008, GO:0045456, GO:0001889, GO:2000045, GO:0060706, GO:0022611, GO:0007552 and GO:0007455) were enriched. Overlap between co-expression modules and differential gene expression (cluster 1) in larvae feeding on *C. cordifolia* showed that gene ontology terms related to polysaccharide catabolism (GO:0000272), regulation of methylation (GO:0006211, GO:0019857 and GO:0051595), lipid metabolism (GO:0006638, GO:0010511, GO:0043651, GO:0046503 and GO:0048017), transport of molecules (GO:0015718, GO:0032507, GO:0070861 and GO:1901571), development and/or cell/organ growth (GO:0002119, GO:0007525, GO:0016049, GO:0021681, GO:0040008, GO:0045463, GO:0050770, GO:0061153, GO:0061888 and GO:1902988), reproduction (GO:0060250, GO:2000241, GO:0090176 and GO:0060278), feeding behavior (GO:0042755), response to endogenous stimuli (GO:0071495), locomotion (GO:0060361), signaling (GO:0035556, GO:0033363, GO:1902948, GO:1902771, GO:0099177, GO:0048017 and GO:0033209) and regulation of transcription/translation (GO:0016070, GO:0043974, GO:0048025, GO:0098781) were enriched.

## Discussion

We compared gene expression patterns of adult females of *Pieris macdunnoughii* when laying eggs and third instar larvae when feeding on *Thlaspi arvense* (a toxic and invasive plant) with that of *Cardamine cordifolia* (a native host plant). Although we saw variation in oviposition choice by the females in our assays, we did not find any differential gene expression between the females preferring different plant species. Our analysis revealed 910 host-plant specific transcripts that were differentially expressed in the larvae between the two treatments. Analysis of plant compound detoxification genes in the larvae indicated that larvae that fed on *C. cordifolia* had higher expression of genes involved in detoxification of specialized plant secondary compounds compared to larvae feeding on *T. arvense*. We additionally found five host-plant specific functional gene modules: two in larvae feeding on *C. cordifolia* and three in larvae feeding on *T. arvense*. We found that the larvae feeding on *C. cordifolia* were enriched for feeding behavior, polysaccharide breakdown, development, and upregulation of nucleic acid synthesis, suggesting favorable hostplant feeding. Conversely, the larvae feeding on *T. arvense* were enriched for response to stress, macromolecule metabolism, apoptosis and immune response indicating signs of impaired feeding. Our results dissect the underlying causes of the evolutionary trap and identify the mechanisms decoupling preference-performance in *P. macdunnoughii* in the presence of a novel resource. Overall, our work provides new insights on the consequences of novel resources on native herbivores and highlight ontogenetic specific stages where adaptation is likely to occur.

### Female responses to *Thlaspi arvense*

*Pieris macdunnoughii* has been experiencing *T. arvense* for at least 95 generations (earliest herbarium record of *T. arvense* in Gunnison Valley) and possibly for 150+ generations (since *T. arvense* was first introduced to the Gunnison Valley) [37,50]. Given that fitness costs of laying eggs on *T. arvense* is high[53], the ability of females to differentiate between the native plants and *T. arvense* should be under strong selection. Previous research has identified sex-linked heritable additive genetic variation in oviposition preference of *P. macdunnoughii* females on *T. arvense* [73]. However, the sex-linked heritability was only significant when tested on whole plants or cut stems, but not on leaf extracts of *T. arvense*, suggesting that differences in glucosinolates between the hostplants were not the primary driver of oviposition preference. These results in combination with our current findings highlight that female oviposition preference is not driven by a few genes/alleles of major effect involved in differentiating chemical signatures between the two hostplants.

Although lepidopteran females use a combination of olfactory, visual, and gustatory cues to lay eggs on suitable hostplants, the ultimate step involves gustatory recognition [19,74]. Gustatory receptors are localized in the forelegs of the females and olfactory and visual receptors are located on the antennae and eyes respectively [75,76]. Thus, differences in oviposition preference should translate to heritable changes underlying genes of the sensory system. Additionally, differences in oviposition preference can be driven by differences in sensitivity to the oviposition stimulants present in the hostplants. Electrophysiological readings from antennae and fore tarsi in various herbivorous insects during oviposition have demonstrated differences in neural excitation activity when females experience various oviposition stimulants and can differentiate between relative concentrations of the stimulants [77–79]. These changes, while heritable, may not result in differences in gene expression. Therefore, *T. arvense* could elicit different neural excitation in females that prefer *T. arvense* over those that prefer *C. cordifolia*, even though both share similar cues. This could result from interactions between the oviposition stimulants and other hostplant specific chemicals that female experience when they first land on the leaf of the hostplant. Future experiments including those that measure the neural excitation of gustatory and antennal receptors in females preferring different hostplants will help disentangle the heritability and differential gene expression results.

### Larval responses to *Thlaspi arvense*

Our analysis revealed contrasting transcriptomic patterns in larvae feeding on *C. cordifolia* compared to *T. arvense*. We found that larvae feeding on *C. cordifolia* exhibited signs of adaptive feeding as expected, while larvae feeding on *T. arvense* showed signs of toxicity and inefficient metabolism of compounds present in *T. arvense*.

Our analysis of genes involved in detoxification highlighted these patterns. The detoxification machinery of insects involves three important phases [80]: phase 1 which includes the cytochrome p450 genes and carboxylesterases are involved in redox reactions of plant secondary metabolites [81,82]. Phase 2 enzymes include glutathione-S-transferase, sulfo-transferases and glucuronosyltransferase. Enzymes in phase 2 act on the byproducts of phase 1 enzymes. In specialist herbivores, the plant compounds are directly acted upon by phase 2 enzymes by circumventing the conjugation step on phase 1 enzymes [83]. Phase 3 enzymes, which include ATP binding cassettes, act on the byproducts of phase 2 enzymes and excrete the toxic compounds out of the cell. In addition to these enzymes, trypsin and insect cuticle proteins are involved in improving digestion efficiency [84].

Larvae that fed on *C. cordifolia* had higher expression of a phase 2 enzyme, GST, compared to those feeding on *T. arvense*. This suggests that *P. macdunnoughii* larvae have evolved efficient mechanisms to detoxify the compounds present in *C. cordifolia*, but they are unable to do the same with *T. arvense*. Another phase 2 enzyme, UDPGT, also showed higher expression in larvae feeding on *C. cordifolia* compared to those feeding on *T. arvense*, although the differences were marginally insignificant (fig 2; p=0.059). In contrast, trypsin was the only enzyme that showed higher expression in larvae feeding on *T. arvense* compared to those on *C. cordifolia*. Thus, the larvae feeding on *T. arvense* were unable to effectively employ the specialized phase 2 enzymes to detoxify the toxic products present in the plant. Consequently, the larvae relied on trypsin, an enzyme that increases digestive efficiency to combat these toxic compounds.

Our analysis of enriched genes during larval feeding provided additional insights into the specific processes underlying differences in performance on both hostplants. The larvae that fed on *C. cordifolia* had genes enriched that suggested successful detoxification of plant compounds and adaptive feeding. For example, genes related to feeding behavior and polysaccharide catabolism were enriched suggesting successful breakdown of complex carbohydrates present in the tissues of *C. cordifolia* [85]. In addition, several genes related to macromolecule metabolism (including lipid, linolic acid, 5-methylcytosine and protein depolymerization) were enriched suggesting that larvae were successfully able to assimilate nutrients present in the native hostplant. Subsequently, genes related to nucleic acid metabolism were enriched suggesting that cellular processes were unaffected when feeding on *C. cordifolia*. Lastly, genes related to humoral immune response were also enriched. Immune cell proliferation is also known to be upregulated in other phytophagous insects when feeding on suitable hostplants [86,87].

We identified several densely connected key genes that were associated with larvae feeding on *C. cordifolia*. Our analysis revealed that N-acetyltransferase domain-containing protein (NAT) and uS12 prolyl 3-hydroxylase (PHD) were the top hub genes in the modules associated with *C. cordifolia* feeding larvae. NAT is involved in transacetylation of acetyl-CoA to arylalkylaminase. NAT knockouts in several insects have indicated that it is involved in melatonin synthesis, aromatic neurotransmitter inactivation and cuticle sclerotization. Mutant insects lacking NAT protein have aberrant pigmentation and improper cuticle formation [88]. PHD proteins possess post-translational activity and are involved in many functions including regulation of oxygen and regulation of collagen stability. Normal expression of PHD proteins is critical for preventing cellular hypoxia and is a critical protein in development [89]. Other genes that were identified as top hub genes were involved in normal cellular processes including cell cycle regulation, DNA/RNA synthesis and chromosome structure/function.

In contrast, larvae that fed on *T. arvense* had genes enriched that indicated poor performance and hostplant induced toxicity in the larvae. For example, genes related to general stress response, oxidative stress response, response to reactive oxygen species and acute inflammatory responses were enriched when larvae fed on *T. arvense*. Insect herbivores experience oxidative stress from the secondary compounds present in their hostplants [90,91]. However, insects adapted to their hostplants counteract the oxidative stress through various mechanisms including efficient detoxification of phase 1 and phase 2 enzymes, employing superoxide dismutase and increased levels of reduced glutathione [92]. Larvae that are not adapted to novel hostplants are unable to efficiently employ the above-mentioned strategies and hence experience high levels of oxidative stress and accumulation of reactive oxygen species (ROS) which can lead to reduced performance and higher mortality. The larvae feeding on *T. arvense* also had genes related to apoptotic processes enriched, further supporting the toxic effects of ROS induced by *T. arvense* on cell death. Additionally, larvae on *T. arvense* had glucose metabolism processes enriched, suggesting that the larvae were breaking down cellular glucose reserves for obtaining nutrition instead of breaking down plant polysaccharides as seen in larvae feeding on *C. cordifolia*. Apart from these, genes related to immune cell differentiation were enriched. Natural killer cell differentiation and interleukin-12 production were particularly enriched in larvae when feeding on *T. arvense*. These responses indicate immune responses to fungal and/or bacterial pathogens in the midgut [93]. Plant defenses are known to interact with gut microbiome and can alter the composition from mutualistic bacteria to pathogenic bacteria, especially when larvae feed on low quality or non-host species by initiating leaky guy syndrome [94,95]. Finally, genes related to metamorphosis including ecdysteroid production, response to ecdysone and morphogenesis were upregulated. Many insects respond to unfavorable conditions such as poor hostplant quality by shortening their developmental period and eclosing with smaller adult body size [96].

In larvae that fed on *T. arvense*, densely connected genes included low density lipoprotein receptor-related protein 4 (LDL 4), J domain-containing protein and C2H2-type domain protein in the functional gene modules associated with *T. arvense*. LDL 4 is involved in the transport of cellular cholesterol and is a crucial component in production of ecdysone, suggesting that larvae feeding on *T. arvense* showed accelerated development [97]. J domain-containing protein is part of the hsp70 chaperone machinery involved in several cellular functions including protein folding, transport of polypeptides across organelles, degrading proteins and coordinating responses to stress [98]. C2H2-type domain protein belongs to the zinc finger domain commonly found in transcription factors and is involved in regulating gene expression. These proteins play an important role in regulating gene expression in response to environmental stress [99]. Together, these hub genes indicate that larvae feeding on *T. arvense* are unable to process the compounds present in the plant and are upregulating genes involved in stress response and accelerated development.

### Implications for preference-performance hypothesis and the outcome of evolutionary trap

Our results demonstrate the decoupling of female oviposition preference and larval performance in the presence of *T. arvense*. The fitness costs related to laying eggs on *T. arvense* is dependent on the fine-grained structure of *T. arvense* in the habitat and its proximity to native host plants [46,53]. *Thlaspi arvense* prefers dry habitats and occurs in disturbed environments [51]. The abundance and range of *T. arvense* could increase due to continued anthropogenic disturbance in native habitats, thus increasing the frequency at which *P. macdunnoughii* females encounter the invasive plant. We here, have shown the lack of transcriptomic differences in females preferring *T. arvense* over *C. cordifolia* for oviposition while larvae exhibit host-plant specific transcriptomic differences during feeding. These results suggest that selection would likely act on larval ability to feed on *T. arvense* rather than females’ ability to avoid laying eggs on the plant. Several cases where repeated error-prone oviposition by females have resulted in larvae incorporating the novel host plant has been documented, including in the related species *Pieris oleracea* and its interaction with the invasive mustard *Alliaria petiolata*. After decades of maladaptation on the plant, *P. oleracea* larvae are now able to develop on *A. petiolata* during its bolting stage but not the rosette stage [38,48]. Indeed, recent population genomic analysis of *P. macdunnoughii* from habitats with and without *T. arvense* has shown signatures of selection in *P. macdunnoughii* in response to *T. arvense* in areas where they cooccur. The loci that were under selection were identified to be involved in genes related to metabolism and larval development, thus providing additional support that selection acts on larval survival [100]. One potential route through which larvae can escape the evolutionary trap would be to survive longer on *T. arvense*. This in turn could help larvae to move to native hostplants in the later instars, resulting in rescue from the evolutionary trap.

## Acknowledgements

We thank M. Olson, H. Walton, E. Wagner, O. Shrestha, and S. McDaniel for assisting with animal rearing. We thank R. Steward for providing advice on differential gene expression analysis. We thank W. Watt, J. Quattro, B. Hollis, P. Andolfatto and M. Asher for proving comments on earlier versions of the manuscript. This work was supported by the University of South Carolina Arts & Sciences to CLB and, the Rocky Mountain Biological Laboratory (RMBL) graduate student fellowship to NR. Field research permits were arranged by RMBL.

## Data availability

The raw reads used in the project are deposited at NCBI under the Bioproject accession id PRJNA1076332. The source code and associated data files are available on Github (https://anonymous.4open.science/r/rna_seq_pieris_preference_performance-5A3E/).

## Notes

https://anonymous.4open.science/r/rna_seq_pieris_preference_performance-5A3E/

### Competing Interest Statement

The authors have declared no competing interest.

## References

1. Ehrlich PR, Raven PH. 1964 Butterflies and Plants: A Study in Coevolution. Evolution 18, 586–608.

2. Thompson JN. 1999 The evolution of species interactions. Science 284, 2116–2118. (doi:10.1126/science.284.5423.2116)

3. Thompson JN. 1999 Specific hypotheses on the geographic mosaic of coevolution. American Naturalist 153. (doi:10.1086/303208)

4. Hardy NB, Otto SP. 2014 Specialization and generalization in the diversification of phytophagous insects: tests of the musical chairs and oscillation hypotheses. Proceedings of the Royal Society B: Biological Sciences 281, 20132960. (doi:10.1098/rspb.2013.2960)

5. Jaenike J. 1990 Host Specialization in Phytophagous Insects. Annual Review of Ecology and Systematics 21, 243–273.

6. Joshi A, Thompson JN. 1995 Trade-offs and the evolution of host specialization. Evol Ecol 9, 82–92. (doi:10.1007/BF01237699)

7. Berdegué M, Reitz SR, Trumble JT. 1998 Host plant selection and development in *Spodoptera exigua*: do mother and offspring know best? Entomologia Experimentalis et Applicata 89, 57–64. (doi:10.1046/j.1570-7458.1998.00381.x)

8. Bossart JL, Scriber JM. 1999 Preference Variation in the Polyphagous Tiger Swallowtail Butterfly (Lepidoptera: Papilionidae). Environmental Entomology 28, 628–637. (doi:10.1093/ee/28.4.628)

9. Ladner DT, Altizer S. 2005 Oviposition preference and larval performance of North American monarch butterflies on four *Asclepias* species. Entomologia Experimentalis et Applicata 116, 9–20. (doi:10.1111/j.1570-7458.2005.00308.x)

10. Jaenike J. 1978 On optimal oviposition behavior in phytophagous insects. Theoretical Population Biology 14, 350–356. (doi:10.1016/0040-5809(78)90012-6)

11. Valladares G, Lawton JH. 1991 Host-Plant Selection in the Holly Leaf-Miner: Does Mother Know Best? Journal of Animal Ecology 60, 227–240. (doi:10.2307/5456)

12. Carrasco D, Larsson MC, Anderson P. 2015 Insect host plant selection in complex environments. Current Opinion in Insect Science 8, 1–7. (doi:10.1016/j.cois.2015.01.014)

13. Larsson S, Ekbom B. 1995 Oviposition Mistakes in Herbivorous Insects: Confusion or a Step Towards a New Host Plant? Oikos 72, 155–160. (doi:10.2307/3546051)

14. García-Robledo C, Horvitz CC. 2012 Parent–offspring conflicts, “optimal bad motherhood” and the “mother knows best” principles in insect herbivores colonizing novel host plants. Ecology and Evolution 2, 1446–1457. (doi:10.1002/ece3.267)

15. Rausher MD. 1979 Larval Habitat Suitability and Oviposition Preference in Three Related Butterflies. Ecology 60, 503–511. (doi:10.2307/1936070)

16. Menacer K, Cortesero AM, Hervé MR. 2021 Challenging the Preference–Performance Hypothesis in an above-belowground insect. Oecologia 197, 179–187. (doi:10.1007/s00442-021-05007-5)

17. Gripenberg S, Mayhew PJ, Parnell M, Roslin T. 2010 A meta-analysis of preference-performance relationships in phytophagous insects. Ecol Lett 13, 383–393. (doi:10.1111/j.1461-0248.2009.01433.x)

18. Trowbridge CD, Todd CD. 2001 Host-Plant Change in Marine Specialist Herbivores: Ascoglossan Sea Slugs on Introduced Macroalgae. Ecological Monographs 71, 219–243.

19. Steward RA, Boggs CL. 2020 Experience may outweigh cue similarity in maintaining a persistent host-plant-based evolutionary trap. Ecological Monographs 90, e01412. (doi:10.1002/ecm.1412)

20. Wiklund C. 1975 The evolutionary relationship between adult oviposition preferences and larval host plant range in *Papilio machaon* L. Oecologia 18, 185–197. (doi:10.1007/BF00345421)

21. Janz N, Nylin S, Wedell N. 1994 Host plant utilization in the comma butterfly: sources of variation and evolutionary implications. Oecologia 99, 132–140. (doi:10.1007/BF00317093)

22. Stefanescu C, Jubany J, Dantart J. 2012 Egg–laying by the butterfly *Iphiclides podalirius* (Lepidoptera, Papilionidae) on alien plants: a broadening of host range or oviposition mistakes? Animal Biodiversity and Conservation 29, 83–90.

23. Nylin S, Janz N. 1993 Oviposition preference and larval performance in *Polygonia c-album* (Lepidoptera: Nymphalidae): the choice between bad and worse. Ecological Entomology 18, 394–398. (doi:10.1111/j.1365-2311.1993.tb01116.x)

24. Nylin S, Bergström A, Janz N. 2000 Butterfly Host Plant Choice in the Face of Possible Confusion. Journal of Insect Behavior 13, 469–482. (doi:10.1023/A:1007839200323)

25. Nylin S, Janz N. 2009 Butterfly host plant range: an example of plasticity as a promoter of speciation? Evol Ecol 23, 137–146. (doi:10.1007/s10682-007-9205-5)

26. Calla B, Noble K, Johnson RM, Walden KKO, Schuler MA, Robertson HM, Berenbaum MR. 2017 Cytochrome P450 diversification and hostplant utilization patterns in specialist and generalist moths: Birth, death and adaptation. Molecular Ecology 26, 6021–6035. (doi:10.1111/mec.14348)

27. Erbilgin N, Ma C, Whitehouse C, Shan B, Najar A, Evenden M. 2014 Chemical similarity between historical and novel host plants promotes range and host expansion of the mountain pine beetle in a naïve host ecosystem. New Phytologist 201, 940–950. (doi:10.1111/nph.12573)

28. Futuyma DJ, Agrawal AA. 2009 Macroevolution and the biological diversity of plants and herbivores. Proceedings of the National Academy of Sciences 106, 18054–18061. (doi:10.1073/pnas.0904106106)

29. Celorio-Mancera M de la P, Wheat CW, Huss M, Vezzi F, Neethiraj R, Reimegård J, Nylin S, Janz N. 2016 Evolutionary history of host use, rather than plant phylogeny, determines gene expression in a generalist butterfly. BMC Evolutionary Biology 16, 59. (doi:10.1186/s12862-016-0627-y)

30. Yoon S, Read Q. 2016 Consequences of exotic host use: impacts on Lepidoptera and a test of the ecological trap hypothesis. Oecologia 181, 985–996. (doi:10.1007/s00442-016-3560-2)

31. Keeler MS, Chew FS. 2008 Escaping an evolutionary trap: preference and performance of a native insect on an exotic invasive host. Oecologia 156, 559–568. (doi:10.1007/s00442-008-1005-2)

32. Casagrande RA, Dacey JE. 2007 Monarch Butterfly Oviposition on Swallow-Worts (*Vincetoxicum spp*.). Environmental Entomology 36, 631–636. (doi:10.1603/0046-225X(2007)36[631:MBOOSV]2.0.CO;2)

33. Steward RA, Fisher LM, Boggs CL. 2019 Pre- and post-ingestive defenses affect larval feeding on a lethal invasive host plant. Entomologia Experimentalis et Applicata 167, 292–305. (doi:10.1111/EEA.12773)

34. Wheat CW, Vogel H, Wittstock U, Braby MF, Underwood D, Mitchell-Olds T. 2007 The genetic basis of a plant–insect coevolutionary key innovation. Proceedings of the National Academy of Sciences 104, 20427–20431. (doi:10.1073/pnas.0706229104)

35. Singer MC. 2021 Preference Provides a Plethora of Problems (Don’t Panic). Annual Review of Entomology 66, 1–22. (doi:10.1146/annurev-ento-022720-061725)

36. Singer MC, Parmesan C. 2019 Butterflies embrace maladaptation and raise fitness in colonizing novel host. Evolutionary Applications 12, 1417–1433. (doi:10.1111/eva.12775)

37. Chew FS. 1977 The Effects of Introduced Mustards (Cruciferae) on Some Native North American Cabbage Butterflies (Lepidoptera: Pieridae). Atala 5, 13–19.

38. Huang XP, Renwick JAA, Chew FS. 1994 Oviposition stimulants and deterrents control acceptance of *Alliaria petiolata* by *Pieris rapae* and *P. napi oleracea*. Chemoecology 5, 79–87. (doi:10.1007/BF01259436)

39. Hale R, Morrongiello JR, Swearer SE. 2016 Evolutionary traps and range shifts in a rapidly changing world. Biology Letters 12, 2016–2019. (doi:10.1098/rsbl.2016.0003)

40. Robertson BA, Chalfoun AD. 2016 Evolutionary traps as keys to understanding behavioral maladapation. Current Opinion in Behavioral Sciences 12, 12–17. (doi:10.1016/j.cobeha.2016.08.007)

41. Robertson BA, Rehage JS, Sih A. 2013 Ecological novelty and the emergence of evolutionary traps. Trends in Ecology and Evolution 28, 552–560. (doi:10.1016/j.tree.2013.04.004)

42. Robertson BA, Hutto RL. 2006 A framework for understanding ecological traps and an evaluation of existing evidence. Ecology 87, 1075–1085. (doi:10.1890/0012-9658(2006)87[1075:AFFUET]2.0.CO;2)

43. Chew FS, Watt WB. 2006 The green-veined white (*Pieris napi* L.), its Pierine relatives, and the systematics dilemmas of divergent character sets (Lepidoptera, Pieridae). Biological Journal of the Linnean Society 88, 413–435. (doi:10.1111/j.1095-8312.2006.00630.x)

44. Chew FS. 1977 Coevolution of Pierid Butterflies and Their Cruciferous Foodplants. II. The Distribution of Eggs on Potential Foodplants. Evolution 31, 568–579. (doi:10.2307/2407522)

45. Chew FS. 1980 Foodplant preferences of Pieris caterpillars (Lepidoptera). Oecologia 46, 347–353. (doi:10.1007/BF00346263)

46. Nakajima M, Boggs CL. 2015 Fine-grained distribution of a non-native resource can alter the population dynamics of a native consumer. PLoS ONE 10, 1–17. (doi:10.1371/journal.pone.0143052)

47. Edger PP et al. 2015 The butterfly plant arms-race escalated by gene and genome duplications. Proceedings of the National Academy of Sciences 112, 8362–8366. (doi:10.1073/pnas.1503926112)

48. Haribal M, Yang Z, Attygalle AB, Renwick JAA, Meinwald J. 2001 A Cyanoallyl Glucoside from *Alliaria petiolata*, as a Feeding Deterrent for Larvae of *Pieris napi oleracea*. J. Nat. Prod. 64, 440–443. (doi:10.1021/np000534d)

49. Haribal M, Renwick JAA. 1998 Isovitexin 6″-O-β-d-glucopyranoside: A feeding deterrent to *Pieris napi oleracea* from *Alliaria petiolata*. Phytochemistry 47, 1237–1240. (doi:10.1016/S0031-9422(97)00740-1)

50. Best KF, McIntyre GI. 1975 The biology of Canadian weeds: 9. *Thlaspi arvense* L. Can. J. Plant Sci. 55, 279–292. (doi:10.4141/cjps75-039)

51. Warwick SI, Francis A, Susko DJ. 2011 The biology of Canadian weeds. 9. Thlaspi arvense L. (updated). https://doi.org/10.4141/P01-159 82, 803–823. (doi:10.4141/P01-159)

52. Rodman JE, Chew FS. 1980 Phytochemical correlates of herbivory in a community of native and naturalized Cruciferae. Biochemical Systematics and Ecology 8, 43–50. (doi:10.1016/0305-1978(80)90024-1)

53. Nakajima M, Boggs CL, Bailey S, Reithel J, Paape T. 2013 Fitness costs of butterfly oviposition on a lethal non-native plant in a mixed native and non-native plant community. Oecologia 172, 823–832.

54. Andrews S. 2010 A quality control tool for high throughput sequence data. See http://www.bioinformatics.babraham.ac.uk/projects/fastqc/.

55. Bolger AM, Lohse M, Usadel B. 2014 Trimmomatic: a flexible trimmer for Illumina sequence data. Bioinformatics 30, 2114–2120. (doi:10.1093/bioinformatics/btu170)

56. Steward RA, Okamura Y, Boggs CL, Vogel H, Wheat CW. 2021 The Genome of the Margined White Butterfly (*Pieris macdunnoughii*): Sex Chromosome Insights and the Power of Polishing with PoolSeq Data. Genome Biology and Evolution 13. (doi:10.1093/GBE/EVAB053)

57. Dobin A, Davis CA, Schlesinger F, Drenkow J, Zaleski C, Jha S, Batut P, Chaisson M, Gingeras TR. 2013 STAR: ultrafast universal RNA-seq aligner. Bioinformatics 29, 15–21. (doi:10.1093/bioinformatics/bts635)

58. Dobin A, Gingeras TR. 2015 Mapping RNA-seq Reads with STAR. Current protocols in bioinformatics 51, 11.14.1--11.14.19. (doi:10.1002/0471250953.bi1114s51)

59. Li H et al. 2009 The Sequence Alignment/Map format and SAMtools. *Bioinformatics (Oxford*, England) 25, 2078–2079. (doi:10.1093/bioinformatics/btp352)

60. Liao Y, Smyth GK, Shi W. 2014 featureCounts: an efficient general purpose program for assigning sequence reads to genomic features. Bioinformatics 30, 923–930. (doi:10.1093/bioinformatics/btt656)

61. Liao Y, Smyth GK, Shi W. 2019 The R package Rsubread is easier, faster, cheaper and better for alignment and quantification of RNA sequencing reads. Nucleic Acids Research 47, e47. (doi:10.1093/nar/gkz114)

62. R Development Core Team. 2008 R: A language and environment for statistical computing.

63. Robinson MD, McCarthy DJ, Smyth GK. 2010 edgeR: a Bioconductor package for differential expression analysis of digital gene expression data. Bioinformatics 26, 139–140. (doi:10.1093/bioinformatics/btp616)

64. Breeschoten T, Ros VID, Schranz ME, Simon S. 2019 An influential meal: host plant dependent transcriptional variation in the beet armyworm, *Spodoptera exigua* (Lepidoptera: Noctuidae). BMC Genomics 20, 845. (doi:10.1186/s12864-019-6081-7)

65. Breeschoten T, Schranz ME, Poelman EH, Simon S. 2022 Family dinner: Transcriptional plasticity of five Noctuidae (Lepidoptera) feeding on three host plant species. Ecology and Evolution 12, e9258. (doi:10.1002/ece3.9258)

66. Kolde R, Kolde MR. 2018 Package ‘pheatmap’. R package 1.

67. Finn RD et al. 2014 Pfam: the protein families database. Nucleic Acids Research 42, D222– D230. (doi:10.1093/nar/gkt1223)

68. Langfelder P, Horvath S. 2008 WGCNA: an R package for weighted correlation network analysis. BMC Bioinformatics 9, 559. (doi:10.1186/1471-2105-9-559)

69. Ritchie ME, Phipson B, Wu D, Hu Y, Law CW, Shi W, Smyth GK. 2015 limma powers differential expression analyses for RNA-sequencing and microarray studies. Nucleic Acids Research 43, e47. (doi:10.1093/nar/gkv007)

70. Huerta-Cepas J, Forslund K, Coelho LP, Szklarczyk D, Jensen LJ, von Mering C, Bork P. 2017 Fast Genome-Wide Functional Annotation through Orthology Assignment by eggNOG-Mapper. Molecular Biology and Evolution 34, 2115–2122. (doi:10.1093/molbev/msx148)

71. Alexa A, Rahnenführer J. 2009 Gene set enrichment analysis with topGO. Bioconductor Improv 27, 1–26.

72. Supek F, Bošnjak M, Škunca N, Šmuc T. 2011 REVIGO Summarizes and Visualizes Long Lists of Gene Ontology Terms. PLOS ONE 6, e21800. (doi:10.1371/journal.pone.0021800)

73. Steward RA, Epanchin-Niell RS, Boggs CL. 2022 Novel host unmasks heritable variation in plant preference within an insect population. Evolution 76, 2634–2648. (doi:10.1111/evo.14608)

74. Pivnick KA, Jarvis BJ, Slater GP. 1994 Identification of olfactory cues used in host-plant finding by diamondback moth, *Plutella xylostella* (Lepidoptera: Plutellidae). J Chem Ecol 20, 1407–1427. (doi:10.1007/BF02059870)

75. McIndoo NE. 1929 Tropisms and sense organs of Lepidoptera. Smithsonian Miscellaneous Collections

76. Xu W. 2020 How do moth and butterfly taste?—Molecular basis of gustatory receptors in Lepidoptera. Insect Science 27, 1148–1157. (doi:10.1111/1744-7917.12718)

77. Den Otter CJ, Schuil HA, Oosten AS-V. 1978 Reception of Host-Plant Odours and Female Sex Pheromone in Adoxophyes Orana (lepidoptera: Tortricidae): Electrophysiology and Morphology. Entomologia Experimentalis et Applicata 24, 570–578. (doi:10.1111/j.1570-7458.1978.tb02818.x)

78. Rojas JC. 1999 Electrophysiological and Behavioral Responses of the Cabbage Moth to Plant Volatiles. J Chem Ecol 25, 1867–1883. (doi:10.1023/A:1020985917202)

79. Beck JJ, Light DM, Gee WS. 2014 Electrophysiological responses of male and female Amyelois transitella antennae to pistachio and almond host plant volatiles. Entomologia Experimentalis et Applicata 153, 217–230. (doi:10.1111/eea.12243)

80. Heckel DG. 2014 Insect Detoxification and Sequestration Strategies. In Annual Plant Reviews, pp. 77–114. John Wiley & Sons, Ltd. (doi:10.1002/9781118829783.ch3)

81. Dermauw W, Van Leeuwen T, Feyereisen R. 2020 Diversity and evolution of the P450 family in arthropods. Insect Biochemistry and Molecular Biology 127, 103490. (doi:10.1016/j.ibmb.2020.103490)

82. Heidel-Fischer HM, Vogel H. 2015 Molecular mechanisms of insect adaptation to plant secondary compounds. Current Opinion in Insect Science 8, 8–14. (doi:10.1016/j.cois.2015.02.004)

83. Engsontia P, Sangket U, Chotigeat W, Satasook C. 2014 Molecular evolution of the odorant and gustatory receptor genes in lepidopteran insects: Implications for their adaptation and speciation. Journal of Molecular Evolution 79, 21–39. (doi:10.1007/s00239-014-9633-0)

84. Dermauw W, Van Leeuwen T. 2014 The ABC gene family in arthropods: Comparative genomics and role in insecticide transport and resistance. Insect Biochemistry and Molecular Biology 45, 89–110. (doi:10.1016/j.ibmb.2013.11.001)

85. Watanabe H, Tokuda G. 2010 Cellulolytic Systems in Insects. Annual Review of Entomology 55, 609–632. (doi:10.1146/annurev-ento-112408-085319)

86. Diamond SE, Kingsolver JG. 2010 Host plant quality, selection history and trade-offs shape the immune responses of *Manduca sexta*. Proceedings of the Royal Society B: Biological Sciences 278, 289–297. (doi:10.1098/rspb.2010.1137)

87. Klemola N, Klemola T, Rantala MJ, Ruuhola T. 2007 Natural host-plant quality affects immune defence of an insect herbivore. Entomologia Experimentalis et Applicata 123, 167–176. (doi:10.1111/j.1570-7458.2007.00533.x)

88. Noh MY, Koo B, Kramer KJ, Muthukrishnan S, Arakane Y. 2016 Arylalkylamine N-acetyltransferase 1 gene (TcAANAT1) is required for cuticle morphology and pigmentation of the adult red flour beetle, Tribolium castaneum. Insect Biochemistry and Molecular Biology 79, 119–129. (doi:10.1016/j.ibmb.2016.10.013)

89. Fong G-H, Takeda K. 2008 Role and regulation of prolyl hydroxylase domain proteins. Cell Death Differ 15, 635–641. (doi:10.1038/cdd.2008.10)

90. Bi JL, Felton GW. 1995 Foliar oxidative stress and insect herbivory: Primary compounds, secondary metabolites, and reactive oxygen species as components of induced resistance. J Chem Ecol 21, 1511–1530. (doi:10.1007/BF02035149)

91. Aucoin R, Guillet G, Murray C, Philogène BJR, Arnason JT. 1995 How do insect herbivores cope with the extreme oxidative stress of phototoxic host plants? Archives of Insect Biochemistry and Physiology 29, 211–226. (doi:10.1002/arch.940290210)

92. Velki M, Kodrík D, Večeřa J, Hackenberger BK, Socha R. 2011 Oxidative stress elicited by insecticides: A role for the adipokinetic hormone. General and Comparative Endocrinology 172, 77–84. (doi:10.1016/j.ygcen.2010.12.009)

93. Shikano I. 2017 Evolutionary Ecology of Multitrophic Interactions between Plants, Insect Herbivores and Entomopathogens. J Chem Ecol 43, 586–598. (doi:10.1007/s10886-017-0850-z)

94. Mason CJ, Ray S, Shikano I, Peiffer M, Jones AG, Luthe DS, Hoover K, Felton GW. 2019 Plant defenses interact with insect enteric bacteria by initiating a leaky gut syndrome. Proceedings of the National Academy of Sciences 116, 15991–15996. (doi:10.1073/pnas.1908748116)

95. Hammer TJ, Bowers MD. 2015 Gut microbes may facilitate insect herbivory of chemically defended plants. Oecologia 179, 1–14. (doi:10.1007/s00442-015-3327-1)

96. Salgado AL, Saastamoinen M. 2019 Developmental stage-dependent response and preference for host plant quality in an insect herbivore. Animal Behaviour 150, 27–38. (doi:10.1016/j.anbehav.2019.01.018)

97. Zhao Y, Liu W, Zhao X, Yu Z, Guo H, Yang Y, Merzendorfer H, Zhu KY, Zhang J. 2023 Low-density lipoprotein receptor-related protein 2 (LRP2) is required for lipid export in the midgut of the migratory locust, Locusta migratoria. Journal of Integrative Agriculture (doi:10.1016/j.jia.2023.07.027)

98. Zhang Y, Liu Y, Guo X, Li Y, Gao H, Guo X, Xu B. 2014 sHsp22.6, an intronless small heat shock protein gene, is involved in stress defence and development in Apis cerana cerana. Insect Biochemistry and Molecular Biology 53, 1–12. (doi:10.1016/j.ibmb.2014.06.007)

99. Guo H, Wang L, Wang C, Guo D, Xu B, Guo X, Li H. 2021 Identification of an Apis cerana zinc finger protein 41 gene and its involvement in the oxidative stress response. Archives of Insect Biochemistry and Physiology 108, e21830. (doi:10.1002/arch.21830)

100. Ravikanthachari N, Steward R, Boggs C. 2023 Local adaptation of a native herbivore to a lethal invasive plant. (doi:10.22541/au.168630489.98488201/v1)

