## Supplementary information for "Genetic Mechanisms underlying Preference-Performance Mismatches: Insights from a Specialized Native Herbivore on an Invasive Toxic Plant"

SI table 1: Paired t-test comparing gene expression (log (CPM)) between *C. cordifolia* and *T. arvense* among different clusters. Adjusted p values reflect BH corrections.

| cluster | . y. | group1 | group2 | n1 | n2 | statistic | df | p | p.adj |
|---|---|---|---|---|---|---|---|---|---|
| 1 | logCPM | *C. cordifolia* | *T. arvense* | 5193 | 5770 | -31.43 | 10750.84 | <0.001 | <0.001 |
| 2 | logCPM | *C. cordifolia* | *T. arvense* | 7047 | 7830 | 41.50 | 14773.26 | <0.001 | <0.001 |
| 3 | logCPM | *C. cordifolia* | *T. arvense* | 1548 | 1720 | -9.51 | 3137.11 | <0.001 | <0.001 |

SI table 2: Paired t-test comparing gene expression (log (CPM)) between *C. cordifolia* and *T. arvense* among different Pfam molecular functions. Adjusted p values reflect BH corrections.

| PFAMs | . y. | group1 | group2 | n1 | n2 | statistic | df | p | p.adj |
|---|---|---|---|---|---|---|---|---|---|
| COesterase | logCPM | *C. cordifolia* | *T. arvense* | 63 | 70 | -1.07 | 131 | 0.288 | 0.288 |
| Cuticle protein | logCPM | *C. cordifolia* | *T. arvense* | 45 | 50 | -1.18 | 82.50 | 0.242 | 0.288 |
| GST | logCPM | *C. cordifolia* | *T. arvense* | 126 | 140 | 4.80 | 258.14 | <0.001 | <0.001 |
| p450 | logCPM | *C. cordifolia* | *T. arvense* | 117 | 130 | -1.86 | 236.16 | 0.0636 | 0.0954 |
| Trypsin | logCPM | *C. cordifolia* | *T. arvense* | 171 | 190 | -3.38 | 342.24 | <0.001 | 0.002 |
| UDPGT | logCPM | *C. cordifolia* | *T. arvense* | 63 | 70 | 2.20 | 115.63 | 0.0295 | 0.059 |

SI table 3: Top 10 hub genes identified by WGCNA analysis for modules that showed significant correlation with feeding on *T. arvense*. Black, blue, and green modules represent positive correlation with *T. arvense* while green and turquoise represent negative correlation with *T. arvense*.

| | | |
|---|---|---|
| **Module blue** | | |
| **Locus id** | **Protein name** | **Function** |
| PMACD_LOCUS16124 | Low density lipoprotein receptor-related protein 4 | transport of cholesterol. Important for ecdysone production. |
| PMACD_LOCUS4367 | Nicotinic acetylcholine receptor alpha 3 subunit | expressed in the nervous system. Important target of insecticides. |
| PMACD_LOCUS4839 | C2 domain-containing protein | calcium-dependent phospholipid binding |
| PMACD_LOCUS3677 | ABC transporter domain-containing protein | Excretion of toxic compounds |
| PMACD_LOCUS6890 | Major facilitator superfamily associated domain-containing protein | membrane transport |
| PMACD_LOCUS2965 | EGF-like domain-containing protein | role in immune response and inflammation |
| PMACD_LOCUS10830 | C2 domain-containing protein | calcium-dependent phospholipid binding |
| PMACD_LOCUS11798 | CUB domain-containing protein | insect serine proteases. Involved in digestion, immunity, and is the main component in many insect venoms |
| PMACD_LOCUS11025 | Protein FAM13A | homolog of this protein in mammals affect fat distribution |
| PMACD_LOCUS4686 | Dipeptidase | expressed in larval mid-gut and is important for digestion. |
| **Module green** | | |
| **Locus id** | **Protein name** | **Function** |
| PMACD_LOCUS12465 | J domain-containing protein | endosome organization |
| PMACD_LOCUS4616 | WAPL domain-containing protein | involved in meiosis |
| PMACD_LOCUS7466 | DEUBAD domain-containing protein | deubiquitinase |
| PMACD_LOCUS12073 | ubiquitinyl hydrolase 1 | Thiol-dependent hydrolysis of ester, thioester, amide, peptide and isopeptide bond |
| PMACD_LOCUS13628 | histone acetyltransferase | regulation of DNA transcription |
| PMACD_LOCUS595 | receptor protein serine/threonine kinase | response to stress |
| PMACD_LOCUS12079 | Ral GTPase-activating protein subunit alpha | GTPase |
| PMACD_LOCUS1886 | Nipped-B protein | regulation of gene expression |

| | | |
|---|---|---|
| PMACD_LOCUS1406 | UDENN domain-containing protein | GTP/GDP exchange activity |
| PMACD_LOCUS6758 | E3 ubiquitin-protein ligase | protein catabolic process |
| **Module black** | | |
| **Locus id** | **Protein name** | **Function** |
| PMACD_LOCUS13892 | C2H2-type domain-containing protein | Zinc finger proteins |
| PMACD_LOCUS12579 | RING-type E3 ubiquitin transferase | metal ion binding |
| PMACD_LOCUS2563 | DNA-directed RNA polymerase subunit beta | DNA transcription |
| PMACD_LOCUS10719 | Rab-GAP TBC domain-containing protein | mitosis regulation |
| PMACD_LOCUS3438 | C2H2-type domain-containing protein | Zinc finger proteins |
| PMACD_LOCUS6109 | BZIP domain-containing protein | regulation of DNA transcription |
| PMACD_LOCUS4227 | DNA polymerase | regulation of DNA transcription |
| PMACD_LOCUS7791 | non-specific serine/threonine protein kinase | autophagy |
| PMACD_LOCUS4415 | C2H2-type domain-containing protein | Zinc finger proteins |
| PMACD_LOCUS8600 | G-protein coupled receptors family 2 profile 2 domain-containing protein | signalling |
| **Module turquoise** | | |
| **Locus id** | **Protein name** | **Function** |
| PMACD_LOCUS7493 | N-acetyltransferase domain-containing protein | sclerotization |
| PMACD_LOCUS2961 | Elongation factor 1-delta | elongation factor involved in protein synthesis |
| PMACD_LOCUS13458 | Large ribosomal subunit protein mL52 | mitochondrial gene expression |
| PMACD_LOCUS12217 | ER membrane protein complex subunit 6 | organelle organization |
| PMACD_LOCUS15646 | Small EDRK-rich factor-like N-terminal domain-containing protein | protein folding |
| PMACD_LOCUS8328 | WKF domain-containing protein | RNA binding |
| PMACD_LOCUS14305 | Exoribonuclease phosphorolytic domain-containing protein | RNA processing |
| PMACD_LOCUS9665 | KRR1 small subunit processome component | Required for 40S ribosome biogenesis |
| PMACD_LOCUS5570 | Ribosomal protein L1 | RNA chaperone activity |
| PMACD_LOCUS12095 | Trafficking protein particle complex subunit 6B | vescicle mediated transport |
| **Module magenta** | | |

| Locus id | Protein name | Function |
|---|---|---|
| PMACD_LOCUS1478 | uS12 prolyl 3-hydroxylase | post-translational activity |
| PMACD_LOCUS13676 | Anaphase-promoting complex subunit 4-like WD40 domain-containing protein | Mitosis regulation |
| PMACD_LOCUS13885 | GH18 domain-containing protein | carbohydrate metabolic process |
| PMACD_LOCUS12675 | Chromo domain-containing protein | chromosome structure/function |
| PMACD_LOCUS3704 | Microtubule-associated protein RP/EB family member 1 | cell division |
| PMACD_LOCUS6776 | Menin | DNA binding |
| PMACD_LOCUS3779 | CWF19-like protein 1 | cell cycle regulation |
| PMACD_LOCUS16298 | Cleavage and polyadenylation specificity factor subunit 1 | nucleic acid binding |
| PMACD_LOCUS1841 | RNA-binding protein cabeza | nucleic acid binding |
| PMACD_LOCUS13496 | Tudor domain-containing protein | RNA metabolism |

SI table 4: GO terms related to biological processes enriched in larvae feeding on *T. arvense*.

| **GO term id** | **Name** |
|---|---|
| GO:0000050 | urea cycle |
| GO:0000082 | G1/S transition of mitotic cell cycle |
| GO:0001779 | natural killer cell differentiation |
| GO:0006091 | generation of precursor metabolites and energy |
| GO:0006546 | glycine catabolic process |
| GO:0007005 | mitochondrion organization |
| GO:0007455 | eye-antennal disc morphogenesis |
| GO:0007552 | metamorphosis |
| GO:0008152 | metabolic process |
| GO:0008213 | protein alkylation |
| GO:0009052 | pentose-phosphate shunt, non-oxidative branch |
| GO:0009249 | protein lipoylation |
| GO:0009585 | red, far-red light phototransduction |
| GO:0009987 | cellular process |
| GO:0010467 | gene expression |
| GO:0016072 | rRNA metabolic process |
| GO:0016570 | histone modification |
| GO:0017004 | cytochrome complex assembly |
| GO:0022611 | dormancy process |
| GO:0022613 | ribonucleoprotein complex biogenesis |
| GO:0030036 | actin cytoskeleton organization |
| GO:0031163 | metallo-sulfur cluster assembly |
| GO:0031346 | positive regulation of cell projection organization |
| GO:0032259 | methylation |
| GO:0032984 | protein-containing complex disassembly |
| GO:0034641 | cellular nitrogen compound metabolic process |
| GO:0039531 | regulation of viral-induced cytoplasmic pattern recognition receptor signaling pathway |
| GO:0042273 | ribosomal large subunit biogenesis |
| GO:0043053 | Dauer entry |
| GO:0043933 | protein-containing complex organization |
| GO:0044237 | cellular metabolic process |
| GO:0045945 | positive regulation of transcription by RNA polymerase III |
| GO:0046847 | filopodium assembly |
| GO:0048844 | artery morphogenesis |
| GO:0060706 | cell differentiation involved in embryonic placenta development |
| GO:0061450 | trophoblast cell migration |
| GO:0097190 | apoptotic signaling pathway |
| GO:0140053 | mitochondrial gene expression |
| GO:1901163 | regulation of trophoblast cell migration |
| GO:1901564 | organonitrogen compound metabolic process |
| GO:2000045 | regulation of G1/S transition of mitotic cell cycle |
| GO:0001889 | liver development |
| GO:0002673 | regulation of acute inflammatory response |

| | |
|---|---|
| GO:0002832 | negative regulation of response to biotic stimulus |
| GO:0006353 | DNA-templated transcription termination |
| GO:0006457 | protein folding |
| GO:0006518 | peptide metabolic process |
| GO:0006839 | mitochondrial transport |
| GO:0006979 | response to oxidative stress |
| GO:0007006 | mitochondrial membrane organization |
| GO:0009262 | deoxyribonucleotide metabolic process |
| GO:0010608 | post-transcriptional regulation of gene expression |
| GO:0018195 | peptidyl-arginine modification |
| GO:0030449 | regulation of complement activation |
| GO:0031099 | regeneration |
| GO:0032695 | negative regulation of interleukin-12 production |
| GO:0032814 | regulation of natural killer cell activation |
| GO:0034614 | cellular response to reactive oxygen species |
| GO:0035041 | sperm DNA decondensation |
| GO:0036336 | dendritic cell migration |
| GO:0043462 | regulation of ATP-dependent activity |
| GO:0043603 | amide metabolic process |
| GO:0044271 | cellular nitrogen compound biosynthetic process |
| GO:0045456 | ecdysteroid biosynthetic process |
| GO:0045476 | nurse cell apoptotic process |
| GO:0045777 | positive regulation of blood pressure |
| GO:0046385 | deoxyribose phosphate biosynthetic process |
| GO:0051131 | chaperone-mediated protein complex assembly |
| GO:0060586 | multicellular organismal-level iron ion homeostasis |
| GO:0061008 | hepaticobiliary system development |
| GO:0070207 | protein homotrimerization |
| GO:0071478 | cellular response to radiation |
| GO:0071826 | protein-RNA complex organization |
| GO:0090069 | regulation of ribosome biogenesis |
| GO:2000232 | regulation of rRNA processing |
| GO:2000257 | regulation of protein activation cascade |
| GO:0001820 | serotonin secretion |
| GO:0006740 | NADPH regeneration |
| GO:0014015 | positive regulation of gliogenesis |
| GO:0019682 | glyceraldehyde-3-phosphate metabolic process |
| GO:0021697 | cerebellar cortex formation |
| GO:0051156 | glucose 6-phosphate metabolic process |
| GO:0051205 | protein insertion into membrane |
| GO:0051792 | medium-chain fatty acid biosynthetic process |
| GO:0070585 | protein localization to mitochondrion |
| GO:0071390 | cellular response to ecdysone |
| GO:1901566 | organonitrogen compound biosynthetic process |
| GO:1904046 | negative regulation of vascular endothelial growth factor production |
| GO:2000243 | positive regulation of reproductive process |

SI table 5: GO terms related to biological processes enriched in larvae feeding on *C. cordifolia*.

| TermID | Name |
|---|---|
| GO:0000272 | polysaccharide catabolic process |
| GO:0001667 | ameboidal-type cell migration |
| GO:0002119 | nematode larval development |
| GO:0003012 | muscle system process |
| GO:0006211 | 5-methylcytosine catabolic process |
| GO:0006638 | neutral lipid metabolic process |
| GO:0006959 | humoral immune response |
| GO:0007156 | homophilic cell adhesion via plasma membrane adhesion molecules |
| GO:0007525 | somatic muscle development |
| GO:0008088 | axo-dendritic transport |
| GO:0008361 | regulation of cell size |
| GO:0010511 | regulation of phosphatidylinositol biosynthetic process |
| GO:0015718 | monocarboxylic acid transport |
| GO:0016049 | cell growth |
| GO:0016070 | RNA metabolic process |
| GO:0019438 | aromatic compound biosynthetic process |
| GO:0019857 | 5-methylcytosine metabolic process |
| GO:0021681 | cerebellar granular layer development |
| GO:0031033 | myosin filament organization |
| GO:0031034 | myosin filament assembly |
| GO:0031647 | regulation of protein stability |
| GO:0032026 | response to magnesium ion |
| GO:0032507 | maintenance of protein location in cell |
| GO:0033209 | tumor necrosis factor-mediated signaling pathway |
| GO:0033363 | secretory granule organization |
| GO:0035556 | intracellular signal transduction |
| GO:0035821 | modulation of process of another organism |
| GO:0040008 | regulation of growth |
| GO:0042755 | eating behavior |
| GO:0043113 | receptor clustering |
| GO:0043170 | macromolecule metabolic process |
| GO:0043519 | regulation of myosin II filament organization |
| GO:0043520 | regulation of myosin II filament assembly |
| GO:0043651 | linoleic acid metabolic process |
| GO:0043974 | histone H3-K27 acetylation |
| GO:0045463 | R8 cell development |
| GO:0046503 | glycerolipid catabolic process |
| GO:0046710 | GDP metabolic process |
| GO:0048017 | inositol lipid-mediated signaling |
| GO:0048025 | negative regulation of mRNA splicing, via spliceosome |
| GO:0048518 | positive regulation of biological process |
| GO:0050434 | positive regulation of viral transcription |
| GO:0050770 | regulation of axonogenesis |

| | |
|---|---|
| GO:0050789 | regulation of biological process |
| GO:0051235 | maintenance of location |
| GO:0051239 | regulation of multicellular organismal process |
| GO:0051261 | protein depolymerization |
| GO:0051595 | response to methylglyoxal |
| GO:0051653 | spindle localization |
| GO:0060250 | germ-line stem-cell niche homeostasis |
| GO:0060278 | regulation of ovulation |
| GO:0060361 | flight |
| GO:0061153 | trachea gland development |
| GO:0061888 | regulation of astrocyte activation |
| GO:0061912 | selective autophagy |
| GO:0070861 | regulation of protein exit from endoplasmic reticulum |
| GO:0071495 | cellular response to endogenous stimulus |
| GO:0090130 | tissue migration |
| GO:0090176 | microtubule cytoskeleton organization involved in establishment of planar polarity |
| GO:0090394 | negative regulation of excitatory postsynaptic potential |
| GO:0098781 | ncRNA transcription |
| GO:0099177 | regulation of trans-synaptic signaling |
| GO:1901571 | fatty acid derivative transport |
| GO:1902771 | positive regulation of choline O-acetyltransferase activity |
| GO:1902948 | negative regulation of tau-protein kinase activity |
| GO:1902988 | neurofibrillary tangle assembly |
| GO:1902996 | regulation of neurofibrillary tangle assembly |
| GO:1904292 | regulation of ERAD pathway |
| GO:2000241 | regulation of reproductive process |

SI figure 1: Sum of square means with different cluster values for identifying distinct clusters in differential gene expression in larval feeding.

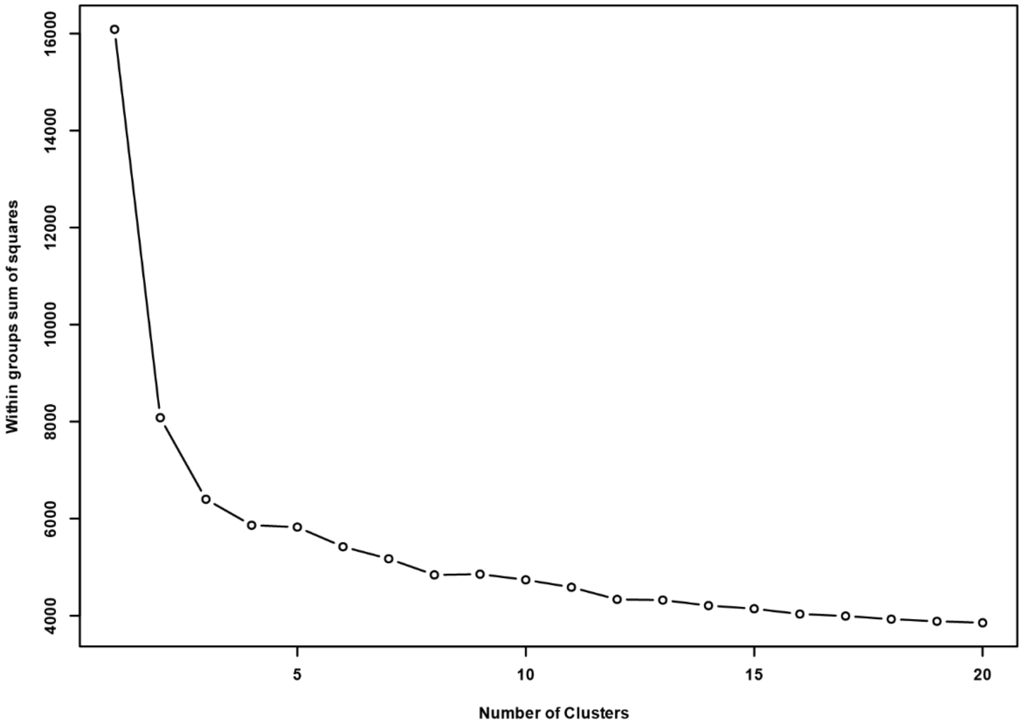

SI figure 2: Scale free independence and mean connectivity values for various soft power values to identify the soft power threshold for constructing signed network of weighted gene co-expression in larval feeding.

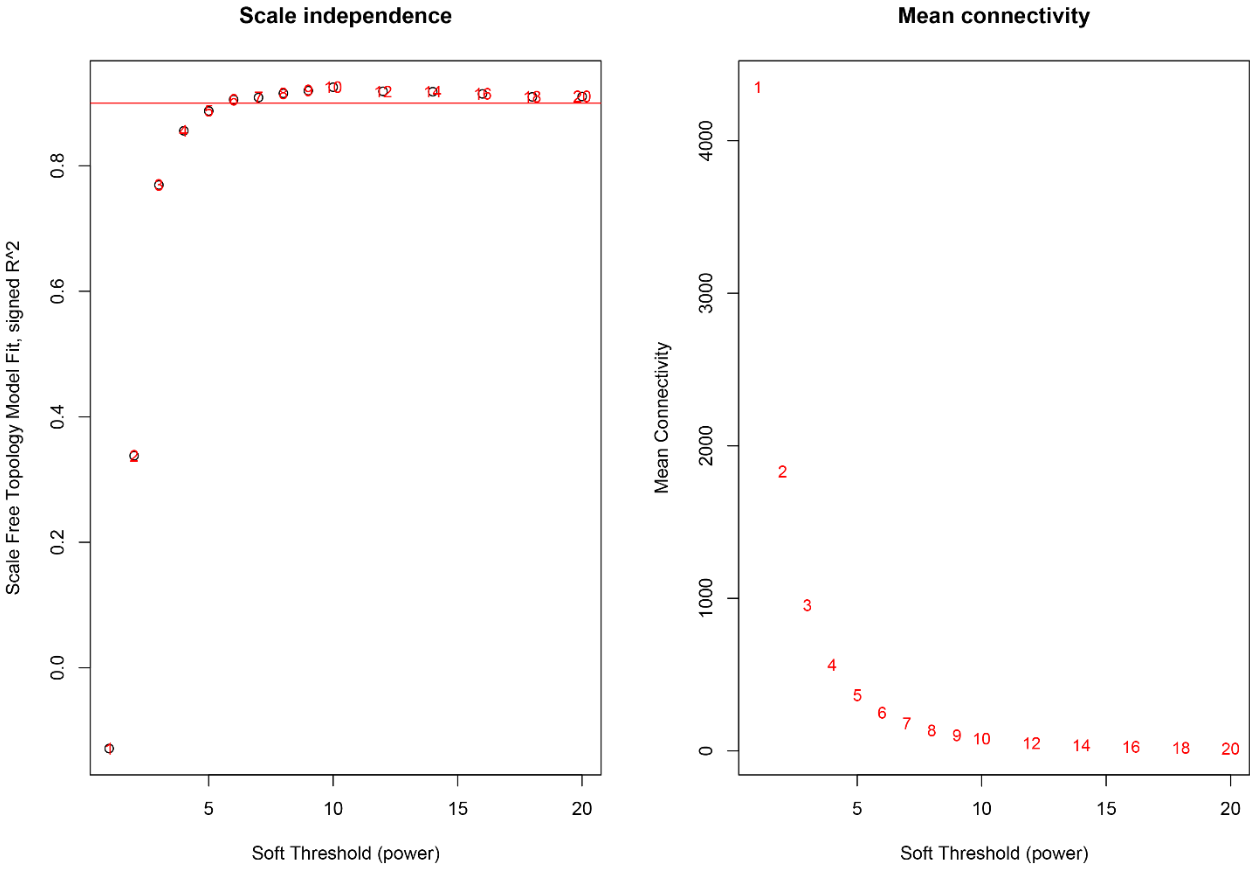

SI figure 3: Dendrogram of different modules (clusters of genes that are co-expressed) identified by the signed network analysis of weighted gene co-expression in larval feeding.

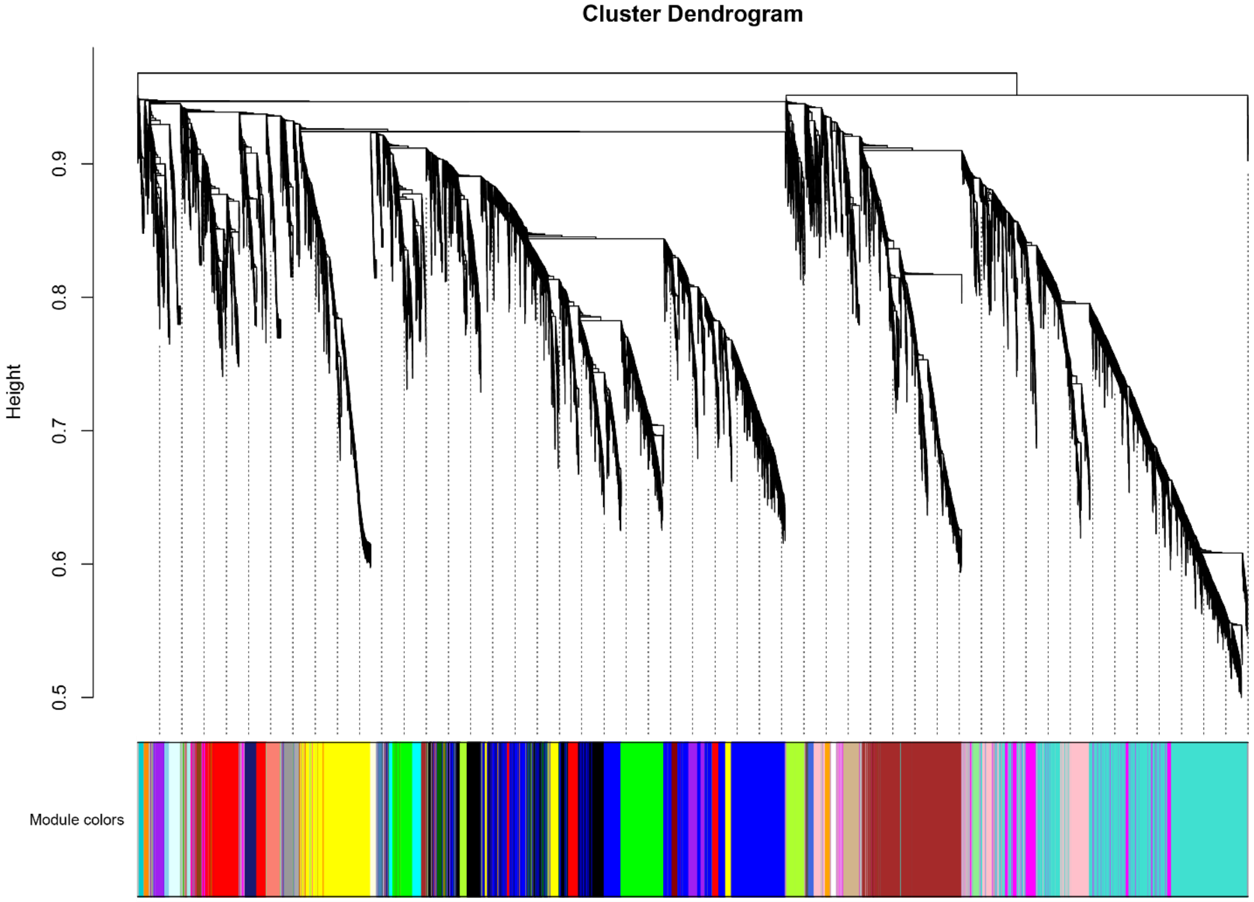
